# Regmex, Motif analysis in ranked lists of sequences

**DOI:** 10.1101/035956

**Authors:** Morten Muhlig Nielsen, Paula Tataru, Tobias Madsen, Asger Hobolth, Jakob Skou Pedersen

## Abstract

Motif analysis has long been an important method to characterize biological functionality and the current growth of sequencing-based genomics experiments further extends its potential. These diverse experiments often generate sequence lists ranked by some functional property. There is therefore a growing need for motif analysis methods that can exploit this coupled data structure and be tailored for specific biological questions. Here, we present a motif analysis tool, Regmex (REGular expression Motif EXplorer), which offers several methods to identify overrepresented motifs in a ranked list of sequences. Regmex uses regular expressions to define motifs or families of motifs and embedded Markov models to calculate exact probabilities for motif observations in sequences. Motif enrichment is optionally evaluated using random walks, Brownian bridges, or modified rank based statistics. These features make Regmex well suited for a range of biological sequence analysis problems related to motif discovery. We demonstrate different usage scenarios including rank correlation of microRNA binding sites co-occurring with a U-rich motif. The method is available as an R package.

## INTRODUCTION

Motif discovery is a classical problem in sequence analysis and its scope broadens with modern sequencing technologies. A large number of tools are designed to find enriched motifs in sequences, with the majority aimed at finding motifs that are enriched in a foreground set of sequences relative to a background set. This is optimal for sequences where a categorical variable defines a foreground and a background. However, many experimental settings are associated with continuous variables where set-based methods are suboptimal. Instead of using a hard threshold to divide a continuous variable into foreground and background, it is more powerful to take the magnitude of the continuous variable directly into account.

More recently, motif enrichment methods have been developed that can exploit the ranking in a list of sequences, e.g. (1, 2, 3, 4, 5). These methods seek to find the motifs that best correlate with the ranked sequence list. Most commonly, this is achieved by exhaustively searching through the space of all simple motifs of a given length k (k-mers). K-mers, ranked by their correlation measures, are then either output directly, clustered and used to define position weight matrices (PWMs) or used as seeds in a variety of downstream algorithms to refine the top correlating motifs.

A general challenge of motif analysis, and specifically of methods based on an exhaustive search, is the rapid increase in search space with motif size and complexity. This problem has been addressed in recent work by using suffix trees, allowing exhaustive searches of large spaces such as all variable gap motifs up to a given length (4). However, functional motifs may display a much higher degree of complexity than current methods meet. Many snoRNAs, for example, are known to bind their targets in a composite motif consisting of two binding sites separated by a variable number of nucleotides. In addition, regulation of biological systems often rely on multiple factors acting in concert. For instance, endogenous RNAs can severely perturb regulatory networks of microRNAs (6). It is thus valuable to be able to evaluate enrichments for subsets of binding sites in combination.

A central aspect in motif analysis of ranked sequences is the significance evaluation of the motif rank correlation. A number of approaches have been used, including linear regression models (7), Wilcoxon rank sum tests (8), a Kolmogorow-Smirnov based approach (9), a Brownian bridge based approach (2) and methods using variants of hyper geometric tests (1, 4, 5). The various methods also have different standards for motif scoring in the sequences. Examples include simple presence/absence scores for each sequence (5, 9), dependence of sequence lengths and global base composition (1) and probabilistic scoring that models base composition of every sequence in the rank list (2).

Presence/absence scores in particular suffer a risk of bias because sequence length and composition is not included in the score model, which is a problem if e.g. sequence lengths are biased in the rank. Also, presence/absence score-based methods may be underpowered in situations where the number of motif occurrences in a single sequence matters.

Based on these issues, we see a need for a tool that allows hypotheses for flexible motifs to be evaluated, and calculates accurate sequence dependent *p*-values for motif observations. We present Regmex, a motif enrichment tool, with a number of new features aimed at accurate significance evaluation (see Figure 1). First, we calculate sequence specific motif *p*-values that depend on both sequence lengths and base compositions using an embedded Markov model. Second, depending on the problem and hypothesis, motif rank correlation or motif clustering can be evaluated in one of three different ways; (1) a Brownian bridge based approach, (2) a modified sum of ranks method which takes sequence properties into account, or (3) a random walk based method which is sensitive to clustering of motif observations anywhere in the sequence list. In addition, Regmex makes use of regular expressions, thus allowing a motif to be far more complex than k-mers. We illustrate some of the benefits by using Regmex on simulated data as well as on real data sets. The method is available as an R package (https://github.com/muhligs/regmex">).

**Figure 1.**
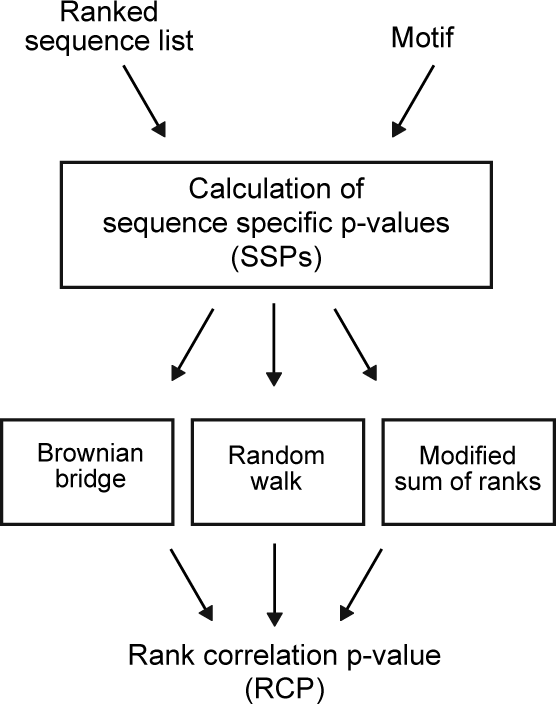
Flow diagram of the procedures for calculating sequence specific *p*-values and rank correlation or clustering p-value in Regmex.

## MATERIALS AND METHODS

### The Regmex tool

In this study, we introduce Regmex, a motif analysis tool available as an R package. Regmex is designed with flexibility in mind to study rank correlation or clustering of motifs in a list of sequences. Briefly, it takes as input a list of sequences ranked by an experimental setting, and one or more motifs, each defined as a regular expression (RE) (see Figure 1). The output, in its simplest form, contains the rank correlation or clustering *p*-values (RCPs) for the input motifs. Alternatively, it is possible to get the underlying sequence specific *p*-values (SSPs) for motifs as well as count statistics etc.

To illustrate the power of REs in a biological sequence context, we consider the following examples:

1. A stem loop structure TTTCNNNGAAA found in the 3’UTR of many key inflammatory and immune genes (10). Although this is a simple RE, it captures 64 11-mers in one expression, and Regmex reports the rank correlation *p*-value of the combined set.
2. A G-quadroplex structure, GGGLGGGLGGGLGGG, 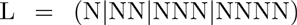. This is found e.g. in telomeric regions (11).
3. Any size open reading frame, 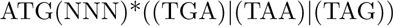. This RE is an example of an enormous set, which would be difficult to obtain without a RE.

We note that an advantage of REs in relation to the motif enrichment problem is that a RE can be obtained for any set of simple motifs. Thus for example a set of experimentally verified binding sequences can be expressed as a RE, and matching will include exactly this set.

### Sequence specific motif *p*-value calculation

A central point in the way Regmex calculates a motif RCP is to calculate SSPs for observing the motif the observed number of times (*n_obs_*) or more. Briefly, from a deterministic finite state automaton (DFA) associated with the RE motif, we identify a sequence specific transition probability matrix (TPM) which is used to build an embedded TPM (eTPM) specific for *n_obs_* (see Figure 2). The SSP is subsequently read from the eTPM raised to the power of the sequence length.

**Figure 2.**
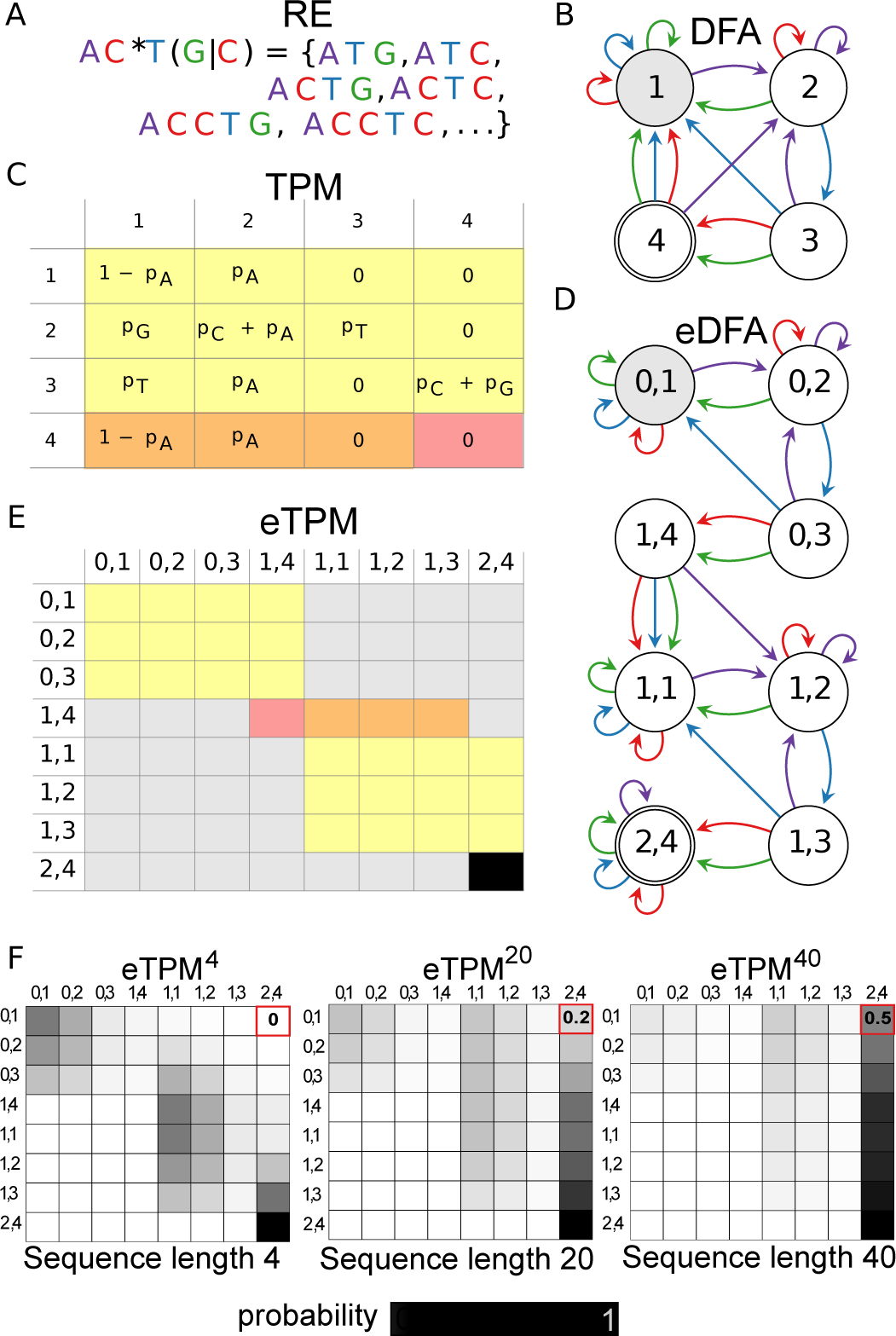
**(A)** Motif in the form of a regular expression. Base coloring applies throughout the figure. **(B)** Deterministic finite state automaton corresponding to the regular expression in (A). Initial state is indicated in gray, end state is indicated by a double circle. **(C)** Transition state probability matrix (TPM) associated with the model in (B). **(D)** Embedded Markov Model (eDFA) for two observed occurrences of the motif. States are pre-indexed with number of prior motif observations. **(E)** Embedded transition state probability matrix (eTPM) associated with the eDFA. The yellow matrix is an exact copy of the yellow matrix from (C). The gray entries have zero probability. The end state transition probabilities of the DFA model (red/orange in (C)) are shifted forward and contain the initial state of the next motif occurrence, except for any end to end transition probability (occurs for REs ending with a *), which remains in the DFA template (red field). The final state of the eDFA (2,4 in D) is an absorbing state and all transition probability is in (2,4;2,4) indicated in black. **(F)** Heat diagrams of the *n*-step eTPM reflecting the probability of moving between states in the eDFA given a random sequence of length *n* with a specific base composition. The row corresponding to the initial state (0,1) holds the probability distribution of going from the start state to any state in the eDFA in *n* steps. The last entry of this row (red field) holds the probability of the observed number of motifs (*n_obs_*) or more in the sequence (the SSP).

#### Deterministic Finite State Automaton

For any RE, the corresponding DFA can be built, which is the initial step in the SSP calculation (Figure 2B). The DFA starts in an initial state, accepts symbols (i.e. nucleotides) on the edges and moves through the states. The end state corresponds to having observed the RE. The DFA used here recognizes an extended regular expression, as described in (12). The routine used to build the DFA for a given RE is implemented in Java, using (13), and supports standard regular expression operations (concatenation, union and Kleene star).

#### Markov Embedding

The DFA graph structure can also be thought of as a Markov model, where instead of accepting symbols, it generates symbols on the edges with probabilities corresponding to the base frequencies in a given sequence. The Markov model can be represented by a transition probability matrix (TPM), which holds the probabilities of moving between states of the DFA given a randomly picked base from the sequence (Figure 2C). TPM*^n^* will hold the probability of moving between states given *n* bases.

We are interested in the SSP and thus need to have a probability model that takes *n_obs_* into account. Regmex does this by making a model expansion using the DFA as a template. We refer to this as an embedded DFA (eDFA) (Figure 2D). Specifically, the template DFA is copied *n_obs_* times and outgoing edges of the end state(s) of the DFA template are moved to the corresponding states in the next template copy. This effectively allows the embedded model to count how many times the RE motif has been observed. The final state of the eDFA is absorbing, so no further motif observations are scored.

Again, the eDFA can be thought of both as an automaton accepting symbols or as a Markov model generating symbols on edges. As above, Regmex constructs a transition probability matrix (eTPM) based on the eDFA (Figure 2E). The eTPM*^n^* holds probabilities of moving between states of the eDFA given a random sequence of length *n* with the observed base frequencies (Figure 2F). We can now extract the probability distribution of the RE motif in a given sequence by reading the row corresponding to the initial state (0,1) in the eTPM*^n^*. In particular the probability of observing the motif *n_obs_* number of times or more (SSP) can be read in the final state column of the initial state row (red field in Figure 2F).

### Motif Rank Correlation *p*-value

In the downstream analysis, Regmex uses the calculated SSPs when calculating the RCP. Because the characteristics of rank correlation may vary depending on the problem analyzed, the choice of method used to evaluate the correlation may differ in detection power. E.g. one test may have higher power for detecting long motifs occurring rarely in the sequence list and another may have superior sensitivity for frequent short motifs. In Regmex, we have implemented three methods for evaluating motif rank correlation or motif clustering, which have different strengths. These methods are based on Brownian bridge (BB), random walk (RW) and modified sum of rank (MSR) statistics. Figure 3 illustrates the concept underlying each of these statistics on a small list of 50 sequences with an enriched motif.

**Figure 3.**
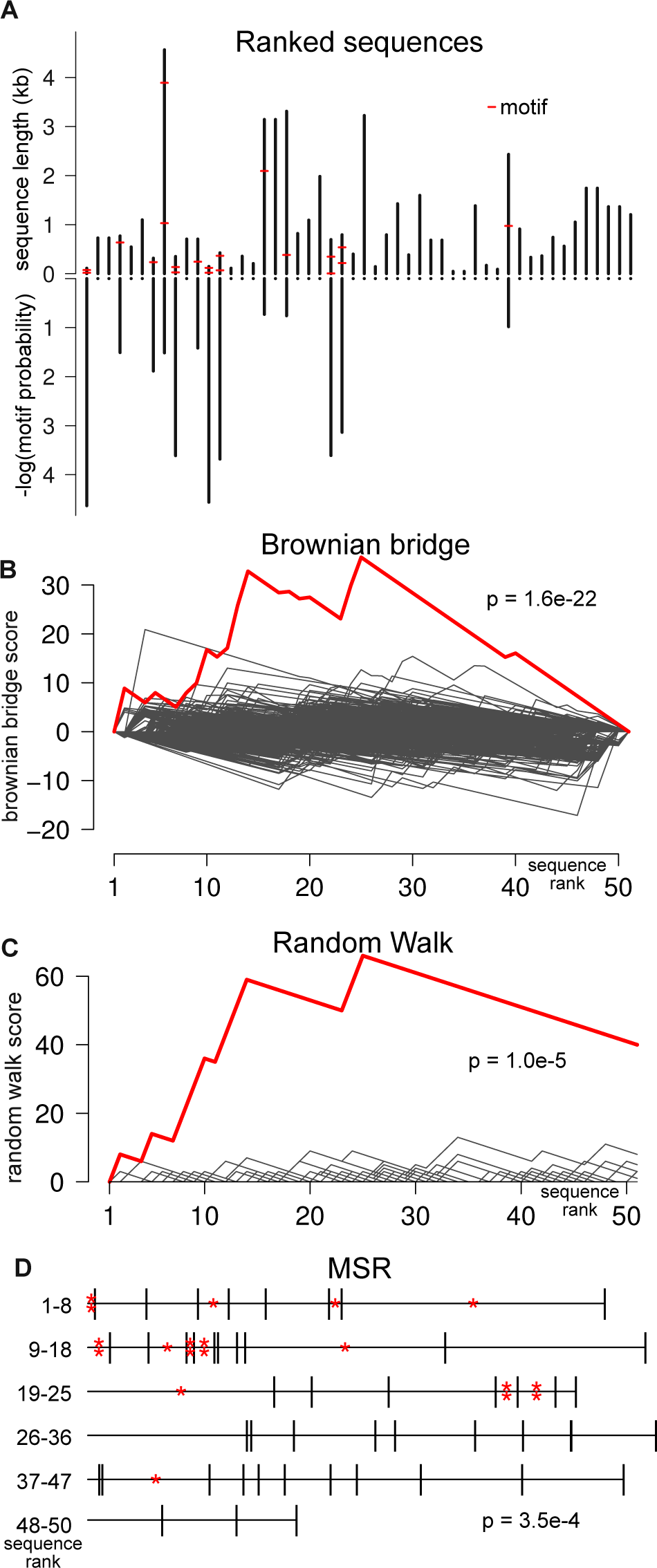
Sequences enriched with a 7-mer motif (ACGTGAT) as indicated with red marks. Upper bars indicate sequence lengths, lower bars indicate SSPs for the motif. **(B)** Brownian bridge for the 7-mer motif in (A) (red) and for 500 random 7-mer motifs (gray). The RCP corresponding to the BB is indicated. **(C)** Random walk for the motif in (A) (red) and 500 random 7-mer motifs (black). The RCP corresponding to the RW is indicated. **(D)** Schematic of the MSR method. Lines represent sequences with lengths proportional to the probability of observing the motif one or more times. A motif occurrence is marked by an asterix. The RCP corresponding to the motif distribution is indicated.

#### Brownian Bridge Method

This method is a re-implementation of the method developed by Jacobsen et al. (14) and recently implemented in cWords (2). Our implementation differs in the calculation of the SSPs and in how we calculate the RCP (see supplemental methods for details). Briefly, the method calculates the max value *D* of a running sum of mean adjusted log scores of the SSPs. The running sum starts and ends in zero and hence is a Brownian bridge under the null model (see Figure 3B). We identify the *p*-value from the analytical distribution of max values of a Brownian bridge.

#### Random Walk Statistic

The RW method is similar to the use of random walks in the BLAST algorithm (15). This method is sensitive to clustering of motifs anywhere in the sequence list. The SSPs for a motif are transformed into steps in a walk (see Figure 3C). Under the null model the motif is not enriched and SSPs follow the uniform distribution. The SSPs are transformed into steps according to a scoring scheme where small *p*-values (SSPs) corresponds to a positive step and large *p*-values corresponds to a negative step. The exact scoring scheme is based on assumed motif densities in the foreground relative to the background, so that higher motif densities give rise to a higher walk in local regions of the sequence list. The RW starts over from zero every time it reaches the lower bound of -1. This makes the RW method sensitive to local runs of enriched motifs in the sequence list. For significance evaluation, we find the probability of a walk with at least as high a max value under the null distribution. We do this using a recursion on an analytic expression for the max value distribution of random walks (see supplemental methods for details). Alternatively, we can use a geometric-like distribution (Gumbel distribution) as an approximation for the max value distribution (16).

#### Modified Sum of Ranks Statistics

The MSR method is based on the idea of using a sum of rank test to determine a rank bias in motif containing sequences. Rather than summing ranks, MSR uses a sum of scores specific for the sequences and motif. The scores are based on sequence specific probabilities, which eliminates bias from sequence composition and length. All motif observations are associated with a score (see supplemental methods for details) that reflect the probability of the motif being found one or more times in the sequence, as well as the rank of the sequence. The score can be considered as a rank normalized for the probability of observing motifs in the sequence. For the MSR method, the null model is that all observed motifs are randomly distributed in the sequences given the sequence compositions and lengths. The test statistic is the sum of the scores, which is approximately normally distributed when the motifs fall randomly in the sequences. The MSR method is faster than the others because we need only the probability of observing one or more motifs in the sequence, which can be read from the TPM of the DFA (Figure 2C) modified so that the end state is absorbing, and thus we do not need to construct the larger embedded model.

## RESULTS

### Combined motifs increase power

Because of different characteristics of the three methods for rank correlation evaluation, they perform differently in different scenarios. Figure 4A illustrates their behavior when applied to a set of 1000 random sequences with a simple 7-mer motif inserted up to 100 times in the upper half of the sequence list. In this particular scenario, the RW approach has the highest sensitivity, followed by the BB method and the MSR method (Figure 4A). The RW method generally has a high sensitivity when the motif density is high, regardless of where in the sequence list it occurs. This is in contrast to both the MSR and BB methods, which are more sensitive to enrichment in the beginning or end of the sequence list. The rank sum derived nature of the MSR method yields a higher sensitivity for enrichment in the ends of longer sequence lists, while the BB method is highly superior in short sequence lists with moderate enrichment. (see Supplemental Figure S1).

**Figure 4.**
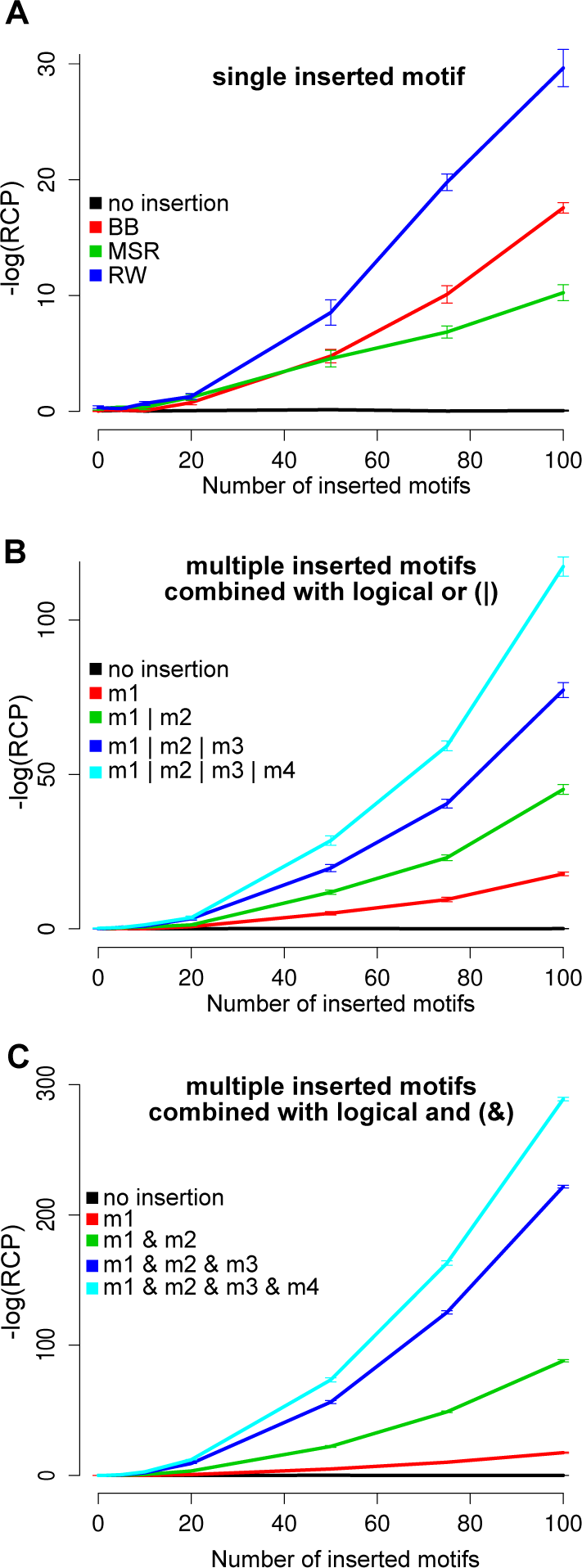
Regmex behavior in different scenarios. **(A)** Comparison of *p*-value output for the different rank correlation methods used in Regmex. One 7-mer motif (ACGTGAT) is inserted as indicated in the first half of 1000 sequences each with a length of 1000 bases. In replicates with no insertion the BB method was used, but the other methods gave similar results. Error bars indicate standard error of 100 replicates. **(B)** Up to four different 7-mer motifs inserted randomly in the first half of the sequences in (A). *p*-value output from Regmex using the BB method plotted against the number of inserted motifs. RE motifs define sets of one up to all four 7-mers as indicated, e.g. 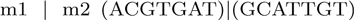. **(C)** *p*-value output from Regmex using the BB method plotted against the number of inserted motifs. Sets of four 7-mer motifs were inserted at fixed positions randomly among the first half of the sequences in (A), so that motifs occur together in the same sequences. RE motifs define sets of combinations of one up to all four 7-mers as indicated, e.g. 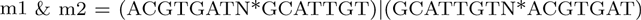, where N denotes any nucleotide.

When using SSPs rather than e.g. binary scores for motif observations in sequences, the benefit of differential scoring becomes clear. This means, e.g., that rank correlation of common and individually insignificant motifs can be better evaluated because their impact on the rank correlation is moderated by the significance of the observation. The same argument applies to rare, highly significant motifs. This, combined with RE motif definition, is useful in the case of evaluating rank correlation of combinations of motifs.

We used Regmex to evaluate rank correlation of combinations of inserted motifs in a set of random sequences. First, we inserted four different simple 7-mers up to 100 times at random positions in the upper half of the ranked sequences. We looked at the behavior of Regmex when defining motifs as REs capturing different subsets of the 7-mers including from one up to all four (i.e. REs defined to capture presence of any member of the subset). We clearly see the effect of combining multiple simple motifs in a set (Figure 4B). When searching for motif 1 or 2 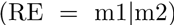, we see a marked increase in detection sensitivity starting at around 20 inserted motifs. As expected, this increases with number of inserted motifs. Rank correlation increases even more dramatically for the motif subsets of three or four 7-mers. We note that the SSPs become less significant when including more 7-mers in the motif, but because the number of inserted motif observations in the enriched end of the sequence list increases (up to 400 for four 7-mers vs. 100 for a single 7-mer), the RCP becomes more significant.

We next looked at the behavior of Regmex when calculating rank correlation of multiple motifs present in the same sequences. Such calculations may be relevant when two or more different factors acting on the same sequences could explain the sequence ranking. To this end, we inserted the four 7-mers together in the same sequences. This was done up to 100 times in the upper half of the sequence list. We used Regmex to calculate RCPs for subsets of combined motifs, i.e. RE motifs designed to capture the presence of one up to all four 7-mers in the same sequence. The SSPs, in contrast to before, now decrease with the number of 7-mers in the RE subset, whereas number of motif observations are identical for all four motifs (by construction). As expected, the detection power of the combined motifs is much higher than that of a single simple motif (Figure 4C).

These experiments show how more complex motifs, such as motif sets, can be captured by REs with great increase in power.

### U rich motifs and microRNA seed target sites as combined motifs

As an example of a scenario where combinations of motifs are relevant, we looked for rank correlation of microRNA seed site targets in combination with a U-rich motif (URM) in a number of microRNA over-expression data sets. URMs are known to bind HuD/ELAVL4 (17) and their presence in 3’UTRs has been shown to correlate with down regulation in several microRNA over¬expression experiments (14). Based on this finding, a model was proposed where URMs augment microRNA induced destabilization of target mRNAs (14). We used Regmex to calculate RCPs for microRNA seed site targets and combinations of the target and URM1 as identified in (14) with sequence UUUUAAA. This was done using 11 different microRNA over-expression data sets (18, 19).

We first calculated RCPs for the microRNA seed site targets. For all data sets, we saw low RCPs for the relevant microRNA seed site target in 3’UTRs, demonstrating correlation between the motif and down-regulated genes (Table 1).

**Table 1.** Rank correlation *p*-values for URM (U) and seed target (S) motifs

| microRNA | Seed Target(S) | $(SN^*U)\vert (UN^*S)$ | $UN^*S$ | $SN^*U$ | Ref. |
| --- | --- | --- | --- | --- | --- |
| miR-7 | $2.6e-03$ | $1.5e-13$ | $2.4e-11$ | $9.6e-05$ | (18) |
| miR-9 | $6.6e-09$ | $1.5e-17$ | $2.6e-19$ | $1.2e-05$ | (18) |
| miR-16 | $1.8e-178$ | $7.3e-147$ | $5.7e-65$ | $9.2e-76$ | (19) |
| miR-106b | $2.5e-99$ | $9.7e-158$ | $4.5e-145$ | $1.8e-58$ | (19) |
| miR-122a | $3.2e-02$ | $4.1e-05$ | $7.6e-04$ | $2.7e-02$ | (18) |
| miR-128a | $6.6e-19$ | $2.2e-48$ | $4.7e-33$ | $8.2e-21$ | (18) |
| miR-132 | $2.4e-08$ | $3.7e-27$ | $3.9e-33$ | $1.2e-07$ | (18) |
| miR-133a | $5.1e-04$ | $9.4e-09$ | $5.7e-06$ | $4.3e-03$ | (18) |
| miR-142 | $2.4e-05$ | $1.4e-13$ | $1.0e-11$ | $6.0e-05$ | (18) |
| miR-148b | $6.3e-09$ | $1.9e-11$ | $3.2e-12$ | $3.4e-04$ | (18) |
| miR-181a | $7.9e-17$ | $3.9e-53$ | $2.4e-46$ | $5.1e-18$ | (18) |

$p$ -values for RE motifs involving URM1 (U) and microRNA seed site targets (S) in different combinations for microRNA over-expression data sets. All $p$ -values were calculated with the BB method. N denotes any nucleotide.

We next calculated RCPs for microRNA seed site targets and URM1 in combination. To this end, we constructed REs of the form 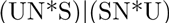, where U denotes the URM, S denotes the microRNA seed site target and N denotes any nucleotide. This RE will capture all combinations of the URM and the seed site in either orientation. As expected, based on the previous findings (14), we consistently saw an even lower RCP for the RE motif capturing both the seed target and the URM1 motif (Table 1). The experiment thus verifies earlier results showing URM1 3’UTR presence correlating with down-regulation.

We next asked whether RCPs are of similar magnitude when the URM is downstream or upstream of the seed target. Here we used Regmex with two REs, SN*U and UN*S for the downstream and upstream question respectively. We observed low RCPs for both the downstream and upstream case for all microRNAs, indicating that URM1 correlates with down-regulation regardless of its relative position to the seed target (Table 1). Notably, we found that RCPs were lower for URM1 upstream the seed target than for URM1 downstream the seed target. This could indicate a preference in relative position between the URM site and the seed target.

The example above illustrates how Regmex is useful in testing well defined hypotheses involving combinations of motifs defined as REs. We note that the outcome verifies earlier findings and further suggests a positional bias in the relative position between the URM motif and the augmented microRNA seed site targets.

## DISCUSSION

We have introduced Regmex, an R package for evaluating rank correlation or clustering of motifs in lists of sequences. Regmex differs from current methods in combining powerful RE motif definition with accurate sequence specific motif significance evaluation and a variety of correlation score statistics. This makes Regmex a flexible tool that expands on the type of motif correlation problems that current methods can handle. Although Regmex handles e.g. traditional exhaustive k-mer screens as other methods (1, 2), it is designed for specific hypothesis testing that involves potentially complex RE motif correlation testing. In particular, Regmex can accurately evaluate rank correlation significance for arbitrary combined sets of simple motifs, such as sets of high scoring k-mers in conventional k-mer screens or combinations of microRNA seed sites. This is relevant for investigations of competitive endogenous RNAs, snoRNA target sites, etc.

The accuracy introduced by the embedded model comes at the cost of computational speed. In particular, motifs that occur frequently in the sequences and are also represented by DFAs with many states require large eTPMs, which may slow down the computation. This can be countered by parallelization, which Regmex natively supports. Furthermore, data structures are simple enough to sub-divide and distribute calculations. Also, the embedded model may be reduced by introducing counting transitions, as suggested in (20).

Regmex offers three alternative ways of evaluating motif rank correlation, which differ in their null models. For the RW method, the null model is that motifs occur at random given the sequence compositions and lengths. The RW method is sensitive to stretches of low SSPs anywhere in the sequence list, and thus may find use in special cases where enrichment is expected off the ends. This could be the case if a sequence list represents consecutive functional sets of sequences, such as a gene ontologies or expression clusters.

Both the MSR and the BB methods are more sensitive to motifs occurring in the ends of the list, but have subtle differences in their null models. For the MSR method, the number of observed motifs in the sequence list is fixed, and only their distribution among the sequences vary under the null. For the BB method, the null is a uniform distribution of SSPs. Although this would suggest a bias for motifs occurring more frequently than expected, the transformation of SSPs into a Brownian bridge via a running sum normalizes for this effect. Thus both of these methods should be robust to motif occurrence bias. As noted (Figure 3 and supplemental Figure S1) they have different sensitivity in different scenarios. The MSR method tends to be more sensitive than the BB method for longer sequence lists and vice versa.

Regmex is implemented in R and offers a number of costumization options including model options such as di-nucleotide dependence and a motif overlap option. Alternative outputs such as SSPs and *n_obs_* combined with simple data formats makes Regmex well suited for a range of problems.

## REFERENCES

1. van Dongen, S., Abreu-Goodger, C., and Enright, A. (2008) Detecting microRNA binding and siRNA off-target effects from expression data. Nature methods, 5(12), 1023–1025.

2. Rasmussen, S. H., Jacobsen, A., and Krogh, A. (2013) cWords - systematic microRNA regulatory motif discovery from mRNA expression data. Silence, 4(1), 2.

3. Steinfeld, I., Navon, R., Ach, R., and Yakhini, Z. (2013) miRNA target enrichment analysis reveals directly active miRNAs in health and disease. Nucleic acids research, 41(3), e45.

4. Leibovich, L., Paz, I., Yakhini, Z., and Mandel-Gutfreund, Y. (2013) DRIMust: a web server for discovering rank imbalanced motifs using suffix trees. Nucleic acids research, 41(Web Server issue), W174–9.

5. Eden, E., Lipson, D., Yogev, S., and Yakhini, Z. (2007) Discovering motifs in ranked lists of DNA sequences. PLoS computational biology, 3(3), e39.

6. Poliseno, L., Salmena, L., Zhang, J., Carver, B., Haveman, W. J., and Pandolfi, P. P. (2010) A coding-independent function of gene and pseudogene mRNAs regulates tumour biology. Nature, 465(7301), 1033–8.

7. Bussemaker, H. J., Li, H., and Siggia, E. D. (2001) Regulatory element detection using correlation with expression. Nature genetics, 27(2), 167–71.

8. Sood, P., Krek, A., Zavolan, M., Macino, G., and Rajewsky, N. (2006) Cell-type-specific signatures of microRNAs on target mRNA expression. Proceedings of the National Academy of Sciences of the United States of America, 103(8), 2746–51.

9. Jensen, L. and Knudsen, S. (2000) Automatic discovery of regulatory patterns in promoter regions based on whole cell expression data and functional annotation. Bioinformatics, 6(4), 326–333.

10. Parker, B. J., Moltke, I., Roth, A., Washietl, S., Wen, J., Kellis, M., Breaker, R., and Pedersen, J. S. (2011) New families of human regulatory RNA structures identified by comparative analysis of vertebrate genomes. Genome research, 21(11), 1929–43.

11. Blackburn, E. H. and Gall, J. G. (1978) A tandemly repeated sequence at the termini of the extrachromosomal ribosomal RNA genes in Tetrahymena. Journal of molecular biology, 120(1), 33–53.

12. Tataru, P., Sand, A., Hobolth, A., Mailund, T., and Pedersen, C. N. S. (2013) Algorithms for hidden markov models restricted to occurrences of regular expressions. Biology, 2(4), 1282–95.

13. Møller, A. dk.brics.automaton – Finite-State Automata and Regular Expressions for Java. (2010) http://www.brics.dk/automaton/

14. Jacobsen, A., Wen, J., Marks, D. S., and Krogh, A. (2010) Signatures of RNA binding proteins globally coupled to effective microRNA target sites. Genome research, 20(8), 1010–9.

15. Altschul, S. F., Gish, W., Miller, W., Myers, E., and Lipman, D. J. (1990) Basic Local Alignment Search Tool. Journal of molecular biology, 215, 403–410.

16. Ewens, W. and Grant, G. (2005) Statistical methods in bioinformatics, Springer Netherlands, 2 edition.

17. Bolognani, F., Contente-Cuomo, T., and Perrone-Bizzozero, N. I. (2010) Novel recognition motifs and biological functions of the RNA-binding protein HuD revealed by genome-wide identification of its targets. Nucleic acids research, 38(1), 117–30.

18. Grimson, A., Farh, K. K.-H., Johnston, W. K., Garrett-Engele, P., Lim, L. P., and Bartel, D. P. (2007) MicroRNA targeting specificity in mammals: determinants beyond seed pairing. Molecular cell, 27(1), 91–105.

19. Linsley, P. S., Schelter, J., Burchard, J., Kibukawa, M., Martin, M. M., Bartz, S. R., Johnson, J. M., Cummins, J. M., Raymond, C. K., Dai, H., Chau, N., Cleary, M., Jackson, A. L., Carleton, M., and Lim, L. (2007) Transcripts targeted by the microRNA-16 family cooperatively regulate cell cycle progression. Molecular and cellular biology, 27(6), 2240–52.

20. Ribeca, P. and Raineri, E. (2008) Faster exact Markovian probability functions for motif occurrences: a DFA-only approach. Bioinformatics (Oxford, England), 24(24), 2839–48.

